# miR-34c-5p is a novel regulator of T cell differentiation that targets FOXP3

**DOI:** 10.1101/2025.07.25.665590

**Authors:** Cláudia Noronha-Estima, Raquel Gaspar-Luta, Tomás Faria Araújo, Carolina Ferreira-Fernandes, Gil Poiares-Oliveira, Margarida Paulo-Pedro, Madalena Pimentel Marques, Ana E Sousa, Margarida Gama-Carvalho

## Abstract

MicroRNAs regulate key genes and pathways essential for T cell differentiation and function; however, many remain poorly characterized. We previously identified miR-34c-5p as a T cell receptor-inducible microRNA in naïve CD4⁺ T cells, but its role in immune regulation remained largely unexplored. In this study, we mapped miR-34c expression across multiple sort-purified CD4⁺ T cell subsets and found that its induction is restricted to FOXP3⁺ cells, both natural regulatory T cells (Tregs) and cells undergoing inducible Treg (iTreg) differentiation trajectory. miR-34c expression correlates with high FOXP3 levels and is absent in conventional memory effector subsets. Functional studies using miR-34c antagomiRs reveal that miR-34c limits iTreg differentiation by restraining FOXP3 expression. Mechanistically, we show that FOXP3 activates miR-34c transcription through direct binding to its promoter, while miR-34c targets the *FOXP3* 3′UTR, establishing a negative feedback loop. Together, our findings identify miR-34c as a FOXP3-responsive miRNA that fine-tunes iTreg development via post-transcriptional repression of *FOXP3*, uncovering a novel layer of regulatory control in T cell responses.

## Introduction

MicroRNAs (miRNAs) are conserved small non-coding RNAs (19-23 nucleotides (nt) long) that regulate gene expression at the post-transcriptional level by targeting mRNAs for degradation or inhibiting their translation in a sequence-specific manner (1). Through this mechanism, miRNAs modulate various aspects of T cell biology, including differentiation, activation, and proliferation (reviewed in (2)). Of note, studies have consistently shown that miRNAs are fundamental for the development and function of T cells, including regulatory T cells (Tregs). For example, mice lacking Dicer, a critical endoribonuclease of the miRNA processing machinery, exhibited impaired T cell development (3,4). Additionally, Dicer deletion specifically in FOXP3⁺ T cells led to fatal autoimmunity resembling the phenotype of FOXP3-deficient mice (5), highlighting the fundamental role of miRNAs in maintaining Treg stability and immune homeostasis.

The regulatory impact of miRNAs is reinforced by their dynamic crosstalk with transcription factors (TFs), forming integrated networks that modulate gene expression at multiple levels. TFs shape the transcriptome by activating or repressing sets of genes, including those encoding miRNAs. In turn, miRNAs post-transcriptionally fine-tune the expression of target mRNAs, often including transcripts encoding TFs. These feedback loops enable precise control over cellular processes such as differentiation, proliferation, and lineage commitment. In negative feedback loops, a TF induces a miRNA that, in turn, represses the expression of this TF, forming a self-limiting circuit that buffers fluctuations and prevents extreme phenotypes (reviewed in (6)). This type of regulation is particularly important in dynamic biological systems such as the immune system. In Tregs, for example, the lineage-defining TF FOXP3 represses miR-31 (7), while miR-31, in turn, directly targets the 3′ untranslated region (UTR) of FOXP3 mRNA to limit its expression (8), forming a negative feedback loop that fine-tunes FOXP3 protein levels and Treg differentiation. Similar circuits have been described for other TF-miRNA combinations, including p53-miR-34a (9) and NF-κB-miR-146a (10), underlining the central role of such feedback loops in ensuring homeostasis and adaptive responsiveness across diverse cellular contexts.

Clarifying the role of miRNAs in immune regulation requires a detailed understanding of T cell molecular regulation, particularly considering the functional heterogeneity of CD4⁺ T cell subsets, each with a distinct role in immune defense and homeostasis. Naïve CD4⁺ T cells serve as precursors of adaptive immune responses, demanding precise regulatory mechanisms to ensure appropriate activation and differentiation, which rely on antigen recognition, co-stimulation, and cytokine signals. These stimuli initiate intracellular pathways that drive transcriptional reprogramming and proliferation of newly activated cells (reviewed in (11)). The cytokine environment then directs their differentiation into specialized subsets, including T helper (Th) cells and Tregs. An environment rich in interleukin-2 (IL-2) and transforming growth factor-beta (TGF-β) is known to promote the differentiation of naïve CD4⁺ T cells into inducible Tregs (iTreg), characterized by FOXP3 and CD25 (IL-2 receptor-α) expression (12), and immune-suppressive properties, vital for maintaining immune homeostasis and preventing autoimmunity (13). Therefore, understanding the molecular regulators of CD4⁺ T cell activation and differentiation is essential for uncovering novel mechanisms that shape immune responses.

We have previously shown that, in addition to the well-characterized miR-155 (14,15), miR-34c-5p (hereafter referred to as miR-34c) is transcriptionally upregulated in naïve CD4⁺ T cells following T cell receptor (TCR) stimulation (16). This observation suggests a potential role for miR-34c in T cell activation and differentiation. Notably, compared to miR-155, miR-34c is undetectable in unstimulated cells and exhibits delayed induction, peaking around four days post-stimulation. In contrast, miR-155 expression peaks two days after stimulation, with levels nearly two orders of magnitude higher than those of miR-34c. While miR-155-5p is one of the most extensively studied miRNAs in the immune system, the biological function of miR-34c in T cells remains unknown.

In this study, we investigate the regulation and functional role of miR-34c in human CD4⁺ T cells. Given the distinct kinetics and magnitude of induction of miR-34c compared to miR-155, we hypothesized that miR-34c may only be expressed in a specific subset of CD4⁺ T cells. To address this, we quantified miR-34c expression in purified CD4⁺ T cell populations, both *ex vivo* and after *in vitro* stimulation. This analysis revealed that miR-34c expression is restricted to FOXP3⁺ cells, prompting us to focus on regulatory T cells and the role of FOXP3 in miR-34c regulation. Using a luciferase reporter assay, we demonstrate that FOXP3 acts as a positive regulator of miR-34c transcription. We then assessed the functional relevance of miR-34c by inhibiting its expression with antagomiRs during naïve T cell activation, followed by transcriptomic analysis and immunophenotyping. Our results show that miR-34c functions as a negative regulator of *FOXP3* expression and constrains the establishment of the iTreg population. Together, these findings identify miR-34c as a novel regulator of Treg differentiation, acting within a FOXP3-centered feedback loop that fine-tunes immune regulation.

## Materials and Methods

### Isolation and culture of human CD4^+^ T cells

Peripheral blood mononuclear cells (PBMCs) were isolated from buffy coats of healthy donors, obtained from *Instituto Português de Sangue e Transplantação* (IPST), by density gradient centrifugation using Lymphosep (Biowest, Cat.L0560 - 500), according to the manufacturer’s instructions. Naïve CD4^+^ T cells were purified from PBMCs using the MojoSort™ Human CD4 Naïve T Cell Isolation Kit (BioLegend, Cat.480042). All purified cell populations were cultured in complete medium (CM, RPMI-1640 + GlutaMAX^TM^ (Gibco, Cat. 61870044) medium supplemented with 10% FBS (Biowest, Cat.S181BH-500), 1% nonessential amino acids (Gibco, Cat.11140050), 1% sodium pyruvate (Gibco, Cat.11360070), 1% penicillin/streptomycin (Gibco, Cat.10378016) and 0.1% gentamycin (Gibco, Cat.15750060)). 1×10^6^ cells/mL of purified CD4^+^ T cells were cultured in 24-well flat bottom plates with CM supplemented with 10 U/mL of human recombinant (hr)-IL-2 (Peprotech, USA, Cat. AF-200-02) and stimulated using CD3/CD28 Dynabeads^TM^ (Gibco, Cat.11131D), at a ratio of 1 bead to 10 cells for 120 h. In specific experiments, the stimulation conditions of naïve CD4^+^ T cells were adjusted to include 5 ng/mL TGF-β (Peprotech, Cat.100-21-50UG) and 100 U/mL of hr-IL-2 (Peprotech, Cat. AF-200-02), to promote the differentiation of FOXP3^+^ cells.

### AntagomiR treatment

miRNA knockdown assays were performed using custom synthesized antagomiR molecules (Horizon Discovery), as before (16). To knockdown miR-34c, freshly isolated naïve CD4^+^ T cells were washed with phosphate buffered saline (PBS) and resuspended in serum-free RPMI 1640 GlutaMAX^TM^ (Gibco, Cat.61870044) to reach a final concentration of 2×10^6^ cells/mL and plated on a 24-well plate. Control or miR-34c-antagomiRs were directly added to the plated cells to reach a final concentration of 1 µM. AntagomiR-treated cells were incubated at 37 °C and 5% CO2 for 2-4 h. After treatment, the medium was replaced, and cells were activated as described above.

### Flow cytometry

Cells were washed once with PBS and surface stained for 25 min at room temperature in the presence of FcR block (Myltenyi, Cat.130-059-901) and fixable viability dye Near-IR (Thermo Fischer Scientific, Cat.L34975), diluted in Brilliant Stain Buffer Plus (BD Biosciences, Cat.566385). Antibodies used for staining are listed as panel 4 in **Table 1**. Then, cells were washed once with PBS, fixed and permeabilized for 30 min at 4 °C using eBioscience™ Foxp3 / Transcription Factor Staining Buffer Set (Invitrogen, Cat.00-5523-00), and, after a wash with permeabilization buffer, intracellularly stained for 30 min at 4 °C. Cells were washed with permeabilization buffer and resuspended in 0.5% BSA in PBS. For RNA-seq experiments only surface staining was performed, using the antibodies listed as panel 3 in **Table 1**. Cells were then fixed with 1% formaldehyde for 10 min before acquisition. Samples were acquired using a 3-laser Cytek Aurora (Cytek Biosciences, USA) spectral flow cytometer. Compensation adjustments after unmixing were performed using the SpectroFlo software (v3.2.1; Cytek Biosciences, USA). Data was analyzed using Flowjo version 10 (BD Biosciences).

Quantitative analysis of flow cytometry data for samples used for RNA-seq was performed using the flowCore (version 2.12.2) (17) and CATALYST (version 1.24.0) (18) Bioconductor packages. Following data import, data for activation markers ICOS, CD25, CD69, CD95, CXCR3, CD45RA, and CD45RO was selected and used to generate the MDS plot using the pbMDS function. Density plot for markers CD45RA and CD69 was generated with the plotExprs function of CATALYST.

### Sorting of CD4^+^ T cell populations

Cells were surface stained as described above using the antibodies listed as panel 1 in **Table 1** and resuspended at a concentration of 50×10^6^ cells/mL in FACS buffer (PBS + 2% FBS + 1 mM EDTA). Naïve and memory Treg, as well as conventional naïve, Th1, Th2 and Th17 CD4^+^ T cells, were sorted according to the strategy described in **Figure 1B**, using a BD FACSymphony S6 SE (BD Biosciences) spectral sorter and collected into FACS tubes containing 0.5 mL of CM. Cells were then lysed for miR-34c quantification or activated for 120 h, as described above. When needed, CD4^+^ T cell enrichment was performed using the MojoSort™ Human CD4 T Cell Isolation Kit (BioLegend, Cat.480010) before cell staining.

**Figure 1.**
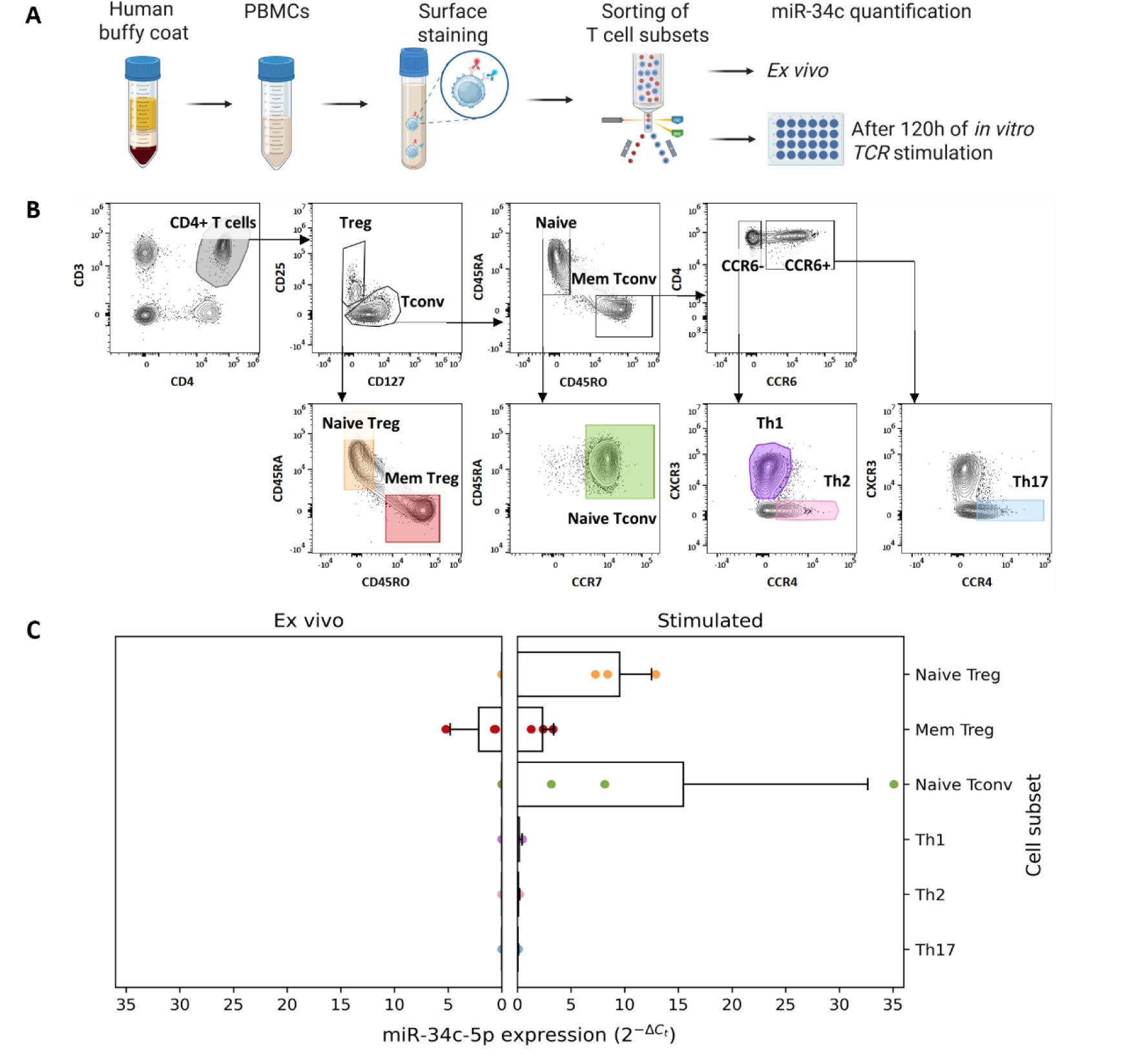
miR-34c expression in diverse human CD4^+^ T cell subsets. **(A)** Experimental workflow for live sorting of T cell populations from human buffy coats, and subsequent mir-34c quantification. **(B)** Sorting strategy to isolate naïve Treg, memory Treg, naïve Tconv, Th1, Th2 and Th17 cell subsets from PBMCs. Color shading represents the gates used for each cell population. **(C)** miR-34c expression levels measured by RT-qPCR in sort-purified cell populations, in both *ex vivo* and *in vitro* stimulated conditions. Data is expressed as mean 2^−ΔCt^ to miR-191-5p ± SD. N = 3 independent donors. N = 3 independent donors.

For sorting of cells for miR-34c quantification in cells with increasing FOXP3 expression the standard staining protocol was modified according to Neyens *et al.* (19), to allow intracellular staining using the antibodies listed as panel 2 in **Table 1**. Briefly, the RNasin® Plus Ribonuclease Inhibitor (Promega, Cat.N2615) was added at 0.4 U/μL during the fixation/permeabilization incubation, at 0.04 U/μL during washes, and at 2 U/μL during the intracellular staining. After the staining, cells were resuspended at a concentration of 50×10^6^ cells/mL in sorting buffer (PBS + 0.1% UltraPure BSA (Ambion, Cat.AM2616) + 1 mM EDTA (Invitrogen, Cat.AM9260G) + 25 mM HEPES (Gibco, Cat.15630080) + 0.4 U/uL RNasin® Plus). Cells with variable FOXP3/CD25 levels were sorted at 4 °C, according to the strategy described in **Figure 2B**, using BD FACSymphony S6 SE (BD Biosciences) and a 70 µm nozzle. Sorted cells were recovered into FACS tubes containing 0.5 mL of sorting buffer. After the sorting, cells were centrifuged at 120xg and resuspended in RNAlater (Invitrogen, Cat.AM7020) in a 1:4 proportion (remaining sorting buffer:RNAlater). Cells in RNAlater were stored at -20 °C until RNA isolation. All sorting experiments were performed at the Flow Cytometry Platform of GIMM - Gulbenkian Institute for Molecular Medicine.

**Figure 2.**
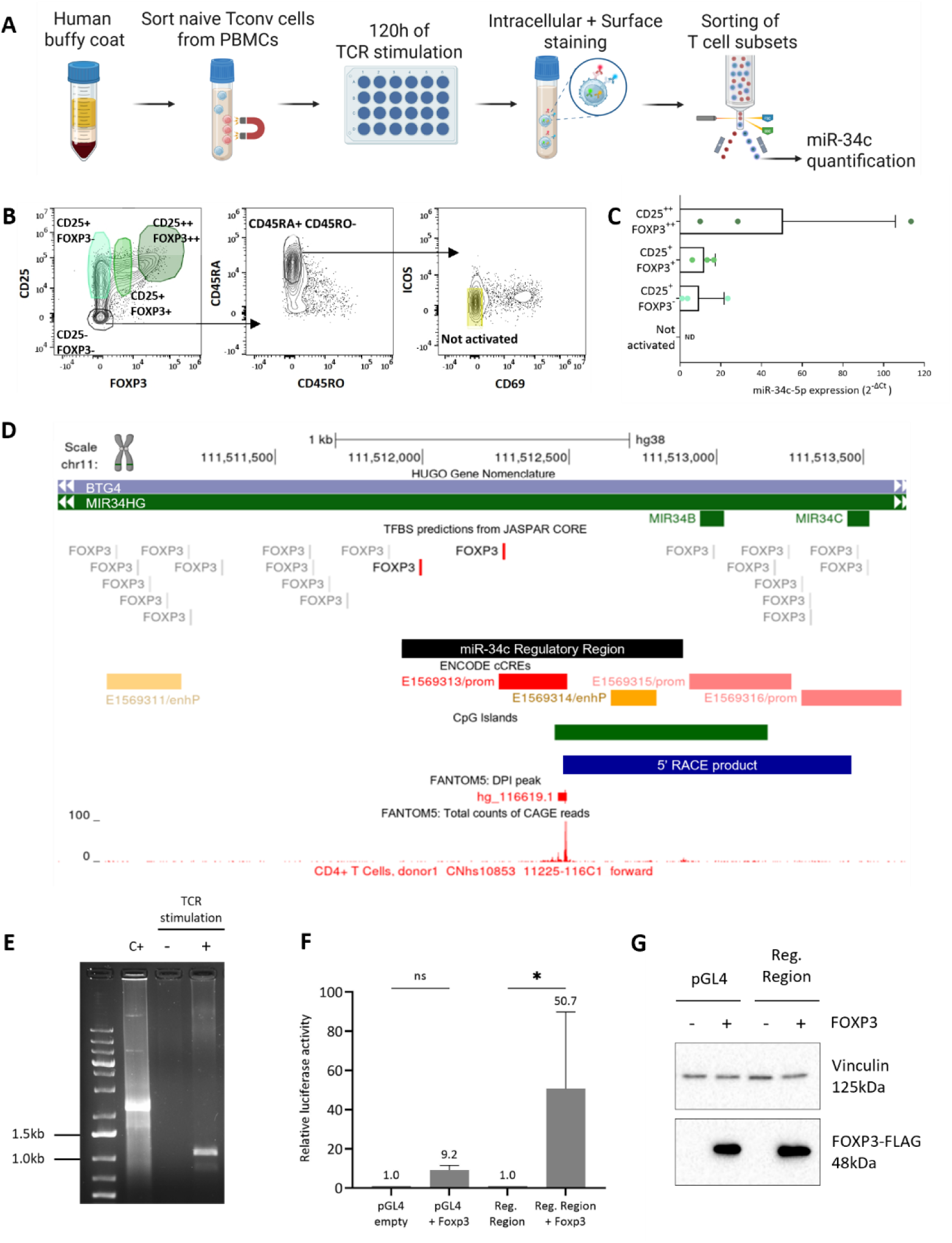
FOXP3 interacts with the miR-34c regulatory region to increase its expression. **(A)** Experimental strategy for the analysis of miR-34c expression levels in *in vitro* stimulated naïve Tconv cell subsets. Stimulated cells were fixed and stained prior to FACS sorting for miR-34c quantification. **(B)** Sorting strategy to isolate stimulated naïve Tconv cell subsets considering their CD25 and FOXP3 expression level. Color shading represents the four selected gates for isolating CD25|FOXP3 high (++), CD25|FOXP3 medium (+) CD25 medium|FOXP3 negative (-) and CD25|FOXP3 negative cells **(C)** miR-34c expression levels measured by RT-qPCR in sorted naïve Tconv cells. Data is expressed as mean 2^−ΔCt^ to miR-191-5p ± SD. N = 3 independent donors. N = 3 independent donors. **(D)** The *MIR34C* genomic *locus* in the GRCh38 assembly, annotated on the UCSC Genome Browser with JASPAR, ENCODE, and FANTOM5 data. Displayed are FOXP3 predicted binding sites (JASPAR CORE), the miR-34c regulatory region cloned for the luciferase reporter assays to analyze its interactions with FOXP3 (black bar), annotated promoter and enhancer regions (red and yellow bars) and CpG island from ENCODE (green bar), and a predicted TSS from FANTOM5 CD4^+^ T cell CAGE datasets. The blue bar represents the mapping of the product amplified by 5’RACE PCR, shown in 2E. **(E)** Agarose gel electrophoresis of the 5’ RACE PCR from total RNA samples of naïve CD4^+^ T cells with (+) and without (-) TCR stimulation. C+ stands for assay positive control with mouse heart total RNA. **(F)** Dual luciferase reporter assay to assess the impact of FOXP3 on the miR-34c regulatory region. HEK-293 cells were co-transfected with the pRL_TK Renilla reporter vector as internal control and a firefly luciferase transcriptional reporter (pGL4 or pGL4-miR-34c regulatory region), in the presence or absence of FOXP3-FLAG over-expression and cell extracts were collected for dual luciferase assays 48 h post-transfection. Plot shows relative firefly to renilla luciferase activity as mean and standard error of the mean for three independent experiments. Data is shown ns – not significant; * - p-value < 0.05 (Mean ± SD. one-way ANOVA) **(G)** Western blot results of FOXP3-FLAG and vinculin (loading control) of a representative sample used for the dual-luciferase assays in panel 2F.

### RNA isolation

Total RNA was isolated from 2×10^5^ - 2×10^6^ human CD4^+^ T cells using miRNeasy Mini Kit (Qiagen, Cat.509217004-K) following the manufacturer’s instructions and eluted in nuclease-free water (NZYTech, Cat.MB04401).

For RNA isolation of sorted cells that underwent fixation and intracellular staining, samples stored in RNAlater were thawed and centrifuged for 30 min at 2500xg at 4 °C to pellet cells. RNAlater was removed, and cells were resuspended in THE RNA Storage Solution (Thermofisher, Cat. AM7000), supplemented with proteinase K at a concentration of 200 ug/mL (Thermofisher, Cat.AM2546). Samples were incubated for 1 h at 60 °C. 4 volumes of TripleXtractor (Grisp, Cat.GB23.0100) were added to each volume of cells in THE RNA Storage Solution. RNA was isolated using the miRNeasy Mini Kit as described above.

### Quantitative reverse transcription polymerase chain reaction (RT-qPCR)

cDNAs for miRNA quantification were synthesized starting from 10 ng of total RNA, using the TaqMan™ Advanced miRNA cDNA Synthesis Kit (Applied Biosystems, Cat.A28007, USA). RT–qPCR assays were performed using the TaqMan™ Fast Advanced Master Mix (Applied Biosystems, Cat.4444557, USA) and the following TaqMan Advanced miRNA Assays (Applied Biosystems, Cat.A25576, USA): hsa-miR-34c-5p (478052_mir), hsa-miR-155-5p (483064_mir) and hsa-miR-191-5p (477952_mir) was used as endogenous control. RT–qPCRs were performed on a CFX96 Touch Real-Time PCR Detection System (Bio-Rad, Cat.1855196, USA) with the following cycling parameters: 95 °C for 20 sec, 40 cycles at 95 °C for 3 sec and 60 °C for 30 sec. All reactions were carried out in 20 μL reaction volumes, in triplicate. The miRNA expression levels were determined by using the 2^-ΔCT^ method.

### miR-34c genomic *locus* analysis

Predicted *FOXP3* Binding sites in the candidate miR-34c promoter region were mapped using the Transcription Factor Binding Sites (TFBS) JASPAR CORE 2020 database (21). Promoter regions and enhancers were identified with ENCODE Candidate *Cis*-Regulatory Elements (cCREs) registry (22,23). H3K4me3 histone modifications profiles were retrieved from public ENCODE Histone ChIP-seq datasets available in the ENCODE portal (24) (https://www.encodeproject.org/). Identification of annotated transcription start sites (TSS) was performed using the FANTOM5 collection (25). All the annotations refer to the Genome Reference Consortium Human Build 38 (GRCh38) and were visualized in the UCSC Genome Browser (26).

### 5’ Rapid Amplification of cDNA Ends (RACE)

5’ RACE was performed using the SMARTer® RACE 5’/3’ Kit (Takara Bio, Cat.634858) according to manufacturer’s instructions. Briefly, 1 µg of total RNA from both unstimulated and stimulated naïve CD4^+^ T cells was directly ligated to the 5’ adapter provided by the kit. Reverse transcription was then carried out using an oligo(dT) primer to generate first-strand cDNA. The 5’-RACE PCR was performed through touchdown PCR using the cDNA as a template and the provided 5’ universal primer (UPM) and the antisense gene-specific primer (**Table 2**), using the following parameters: 5 cycles: 94 °C for 30 sec and 72 °C for 3 min; 5 cycles: 94 °C for 30 sec, 70 °C for 30 sec and 72 °C for 3 min; 30 cycles: 94 °C for 30 sec, 65 °C for 30 sec and 72 °C for 3 min. The 5’-RACE PCR reaction was resolved on a 1% agarose gel from which the DNA was purified. Gel extraction was performed using the NucleoSpin Gel and PCR Clean-Up Kit (Macherey-Nagel, Cat.740609.250). The 5’-RACE PCR products were further amplified by PCR using Platinum™ SuperFi™ II PCR Master Mix (Invitrogen, Cat.12368010), following the manufacturer’s instructions, and validated through Sanger sequencing through STAB VIDA.

### RNA-seq library preparation

Total RNA samples were isolated from purified naïve CD4^+^ T cells from buffy coats of three healthy donors. Cells were cultured in CM and stimulated with CD3/CD28 Dynabeads^TM^ (Gibco, Cat.11131D) at a ratio of 1 bead to 10 cells and 10 U/mL of hr-IL-2 (Peprotech, Cat. AF-200-02). After 40 h, stimulation was removed, cells were treated with either miR-34c or scrambled antagomiRs and cultured for another 100 h. Cells were then harvested, and total RNA was isolated as described above. Purified RNA was quantified using Nanodrop^TM^ One and a minimum of 500 ng of RNA per sample was sent for library preparation and sequencing at the Genomics Unit from Centre for Genomic Regulation (CRG – Barcelona, Spain). Prior to library preparation, all samples underwent quality control using a 2100 Bioanalyzer and the Eukaryote Total RNA Nano Bioanalyzer assay (both from Agilent) to assess RNA integrity (RIN >8) and concentration. Library preparation was performed using poly-A selection. RNA-sequencing was performed using Illumina NextSeq 2000 (2× 50-bp reads, a P2 flowcell and a 100 cycles kit) and a sequencing depth of at least 65 million reads per sample. RNA-seq data is archived under the ENA accession number PRJEB87428.

### RNA-Seq data analysis

Following quality assessment using FastQC version 0.11.5 (https://www.bioinformatics.babraham.ac.uk/projects/fastqc/), Cutadapt (27) was used to remove sequencing adapters and trim the first 10 nt. The trimmed data was filtered using inhouse scripts to remove reads with unknown nt, homopolymers with length ≥ 25 nt or an average Phred score < 30 (28). The remaining reads were aligned to the GRCh38 assembly using the STAR aligner (29) version 2.5.0 with the following options: --outFilterType BySJout -- outFilterMultimapNmax 1 --alignSJoverhangMin 8 --alignSJDBoverhangMin 5 -- outFilterMismatchNmax 1 --alignIntronMax 100000 --outSAMmultNmax 1 --outSAMattributes All --outSAMstrandField intronMotif --outSAMtype BAM Unsorted SortedByCoordinate -- quantMode TranscriptomeSAM GeneCounts --outSAMunmapped Within --outReadsUnmapped Fastx --twopassMode Basic --outSJfilterCountTotalMin 8 5 5 5 --outSJfilterCountUniqueMin -1 -1 -1 -1 --outWigType wiggle read1_5p --limitBAMsortRAM 10000000000. Approximately 90% of the filtered reads were mapped to a single genomic location, corresponding to an average of ∼26 000 detected genes per library. Gene counts were determined using the htseq-count function from HTseq (30) (version 0.9.1) in union mode and discarding low-quality score alignments (–a 10), using the Ensembl GRCh38.98 genome annotation. Data S1 presents the summary information of the RNA-seq datasets. All subsequent data analysis was performed in R (version 4.3.1). Principal Component Analysis (PCA) was performed on normalized gene expression data using DESeq2 (31) (version 1.42.1). Raw counts were transformed with the variance stabilizing transformation after filtering out genes with fewer than 10 counts. PCA was visualized using the plotPCA function, grouping samples by treatment and donor based on the first two principal components. Differential gene expression (DGE) analysis for RNA-Seq gene counts was performed with the limma Bioconductor package (version 3.58.1) using the voom method to convert the read-counts to log2-CPM, with associated weights, for linear modelling (32,33). The design formula included both treatment and donor variables. Samples from each donor were treated as biological replicates, performing a paired analysis to account for inter-donor variability as the donor condition exerted a more substantial influence on sample behavior compared to the treatment condition. Differential expression analysis was conducted using the eBayes function and genes with a |log2foldchange| ≥ 0.6 and p-value ≤ 0.05 were considered differentially expressed genes (DEGs) (the complete list of DEGs can be found in **Data S2**). The EnchancedVolcano (version 1.18.0) (34) and the pheatmap (version 1.0.12) (35) R packages were used to generated the Volcano plot and the heatmap visualization of DEGs, respectively. The code used for analysis is available at https://github.com/GamaPintoLab/miR-34c-RNAseq.

### Gene Ontology (GO) enrichment analysis

Functional enrichment of DEGs was performed with the clusterProfiler package (version 4.10.1) (36–39) using the *Homo sapiens* annotation package “org.Hs.eg.db” in R (version 4.3.1). The false discovery rate of the enriched GO terms was accounted for with the Benjamini-Hochberg method (adjusted p-value cutoff = 0.05). Ten non-redundant GO terms were selected from the top 20 most significantly enriched to generate a lollipop plot for visualization using the ggplot2 (40) R package.

### Plasmids and cloning

The miR-34c regulatory region chr11:111,511,926–111,512,877 (forward strand) was cloned upstream of the firefly luciferase gene in the pGL4.10[*luc2*]-promoterless vector (Promega, E6651). Briefly, it was amplified by slowdown PCR as previously described (41), using 25 ng of Jurkat genomic DNA as input and the primers listed in **Table 2**. The resulting PCR product was amplified to incorporate XhoI and HindIII restriction sites. The insert and pGL4.10[*luc2*] were digested using the same enzymes and ligated using T4 DNA Ligase (Thermo Fisher Scientific, Cat.EL0011) at 22 °C for 1 h, with a 5:1 molar ratio of insert to vector.

The 3′UTR of the *FOXP3* gene, containing the putative miR-34c response sites ChrX: 49,250,575-49,251,318 (GRCh38), was amplified by PCR from genomic DNA isolated from Jurkat cells using the primers listed in **Table 2**. The PCR product was cloned into the pEZX-MT01 plasmid (Genecopoeia, Cat.LPFR-M010) using XbaI and XhoI restriction sites. The pEZX-*FOXP3*-3′UTR-mut plasmid, containing mutated miR-34c MREs, was synthesized by Genscript according to the scheme in **Figure 5A**, by replacing the red-highlighted cytosines for adenines, according to a previous study showing that mismatches in these seed positions abrogate miRNA function (42).

### Western blotting

3×10^5^ HEK-293 cells were lysed in TripleXtractor reagent (Grisp, Cat.GB23.0100) and phase separation was performed following the manufacturer’s instructions. The organic phase was used for protein extraction as described (43).

Protein lysates were loaded onto 10% polyacrylamide gels. After electrophoresis, blotting on a PVDF membrane (Merck Millipore, Cat.IPVH00010) was performed using a wet transfer system and a methanol-based transfer buffer (25 mM Tris, 192 mM glycine, 20% methanol). Blocking was performed with 5% milk in 1x PBST (Phosphate Buffered Saline containing 0.1% (v/v) Tween-20) for 60 min at room temperature with gentle shaking. Blots were incubated with anti-FLAG (1:1000, clone M2, Sigma-Aldrich, Cat.F1804) and anti-vinculin (1:1000, clone 7F9, Santa Cruz Biotechnology, Cat.sc-73614) primary antibodies overnight at 4 °C with gentle agitation, followed by 3x 5 min washes with PBST and a 1 h incubation at room temperature with a goat anti-mouse HRP-conjugated secondary antibody (1:3000 dilution, Bio-Rad, Cat.170-6516). Blots were developed using Clarity^TM^ Western ECL Substrate (Bio-Rad, Cat.170-5060) and immediately analyzed with ChemiDoc XRS+ System (Bio-Rad, Cat.170-8265).

### Luciferase assays

Luciferase assays were performed in HEK-293 cells (ATCC, Cat.CRL-1573), cultured in Minimum Essential Medium (MEM) (Corning, Cat.10-009-CV) supplemented with 10% Fetal Bovine Serum (Biowest, Cat.S181BH-500). Cells were seeded at a density of 1.2 x 10^4^ cells/well on 96-well plates and transfected on the following day using the JetOptimus transfection reagent (Polyplus, Cat.101000051) with a total of 130 ng of plasmid DNA. In parallel, cells were seeded at a density of 6 x 10^4^ cells/well on 24-well plates and transfected under identical conditions using the same plasmid ratio for Western blot analysis to validate FLAG-FOXP3 expression.

For transcription factor reporter assays, cells were transfected with 70 ng of the reporter vector pGL4.10[*luc2*]-promoterless (Promega, Cat.E6651), or pGL4.10[*luc2*]-miR-34c regulatory region, 10 ng of pRL_TK (Promega, Cat.E2241) and 70 ng of expression vector FLAG-FOXP3 (Addgene, Cat.153147) or pBluescriptSK (-) (Stratagene, discontinued). For microRNA reporter assays, cells were transfected with the previously generated miR-34c-pCDNA3.1 construct (16) and either pEZX-MT01-*FOXP3* 3’UTR or pEZX-MT01-empty vector (GeneCopoeia, Cat.LPFR-M010), following a mass ratio of 1 (reporter vector) to 4 (miR-34c expression vector). After 48 h of transfection, culture medium was removed and replaced with 50 µL of MEM per well. Luminescence of Firefly and Renilla luciferase were sequentially measured with a Dual-Glo® Luciferase Assay System (Promega, Cat.E2920) following the manufacturer’s instructions, using the GloMax® 96 Microplate Luminometer (Promega). The mean values of the luciferase to Renilla firefly ratio from three independent assays were compared using two-tailed t-test.

### Statistical analyses

Analyses were performed using the statistical software GraphPad Prism 9.0.0. Data is presented as mean values ± SD. Results were considered significantly different when the p-value was less than 0.05 (*p < 0.05, **p < 0.01, ***p < 0.001, ****p < 0.0001).

### Data availability

The 6 raw fastq files of the RNA-seq data generated for this study have been submitted to the European Nucleotide Archive with the individual study accession number PRJEB87428. All the code used in this work is available at https://github.com/GamaPintoLab/fastfilter and https://github.com/GamaPintoLab/RNA-Seq.

## Results

### High expression levels of miR-34c are exclusively detected in FOXP3^++^ CD4^+^ T cells

We have previously reported the transcriptional upregulation of miR-34c in response to TCR stimulation of conventional naïve CD4^+^ T cells (naïve Tconv, CD4^+^ CD3^+^ CD45RA^+^ CD45RO^-^ CD25^-^) (16). Although miR-34c showed a strong upregulation upon TCR stimulation, its peak expression level was 100 times lower when compared to TCR-induction of miR-155, a well-known regulator of T cell activation. This observation led us to hypothesize that, unlike miR-155, miR-34c expression may be restricted to a specific CD4^+^ T cell subset. To better understand the functional role of miR-34c in the context of T cell activation it is essential to identify the cell types where it is expressed. We therefore sort-purified six main CD4^+^ T cell subsets, including naïve and memory Treg, naïve Tconv, Th1, Th2 and Th17 cells, using an adaptation of a previously described strategy (44). Each subset was sorted and split into two groups for either immediate quantification of *ex vivo* miR-34c levels or *in vitro* stimulation with anti-CD3/CD28 and IL-2 for 120 hours to induce TCR signalling and cell activation prior to miR-34c quantification (**Figure 1A**). Naïve and memory Tregs were included as key regulatory populations, while Th1, Th2, and Th17 subsets represent polarized conventional memory effector states (**Figure 1B**). This approach allowed us to map miR-34c expression across the most distinct and functionally divergent CD4⁺ T cell populations. Results showed that only memory Tregs express detectable levels of miR-34c *ex vivo* (**Figure 1C**). Following *in vitro* TCR stimulation, miR-34c expression was exclusively induced in naïve Treg and naïve T conv cells, while memory Tregs maintained the same expression level (**Figure 1C**). Although all cell types responded to TCR stimulation, as attested by miR-155 upregulation (**Supplementary Figure 1**), miR-34c expression was not detected in any of the stimulated memory T conv subsets (Th1, Th2, Th17). The finding that memory Tregs were the only differentiated cells expressing this miRNA *ex vivo*, along with induction of miR-34c expression in both naïve Treg and naïve Tconv cells in response to TCR stimulation, led us to hypothesize that miR-34c induction could be related to FOXP3 expression - a transient feature of CD4^+^ T cell activation (45) that becomes sustained in thymic/natural Tregs and in naïve T cells cultured under conditions promoting their differentiation into iTregs (46).

To determine whether miR-34c levels are directly associated with FOXP3 expression, we implemented a workflow in which four naïve-derived cell subsets with distinct FOXP3 and CD25 expression profiles were isolated following 120 hours of TCR stimulation (**Figure 2A**) using a previously established protocol that preserves RNA integrity after intracellular staining and fixation (19). We sorted and quantified miR-34c in cells that did not activate (CD25⁻FOXP3⁻CD45RA^+^ICOS⁻CD69⁻), activated cells that did not upregulate *FOXP3* (CD25^+^FOXP3⁻), activated cells that upregulated *FOXP3* (CD25^+^FOXP3^+^) and cells with high expression for both markers (CD25^++^FOXP3^++^) (**Figure 2B**). Although miR-34c was not detectable in cells that did not respond to stimulation, both CD25^+^FOXP3⁻ and CD25^+^FOXP3^+^ cells showed a similar, low expression level of the miR. Of note, the mean expression of miR-34c was three times higher in cells with strong FOXP3 expression (CD25^++^FOXP3^++^) (**Figure 2C**). This pattern was identical in cell populations with comparable levels of CD25/FOXP3 subjected to TCR stimulation in the presence of TGF-β (**Supplementary Figure 2**). These results indicate that the robust upregulation of miR-34c following TCR stimulation is not only a direct consequence of cell activation but also seems to be associated with high FOXP3 expression levels. Since FOXP3 is a transcription factor, we next asked whether it can directly control miR-34c expression.

### FOXP3 is a direct regulator of miR-34c expression

To investigate whether FOXP3 regulates miR-34c transcription, we first retrieved functional annotations related to transcription initiation, including promoter regions and CpG islands, from ENCODE (22). We then identified a potential transcription start site (TSS) in primary CD4⁺ T cells based ion FANTOM5 CAGE (Cap Analysis of Gene Expression) data (25), visualized in the UCSC Genome Browser. In this region, we also identified multiple predicted FOXP3 binding sites using the JASPAR database (**Figure 2D**).

To experimentally validate the reported miR-34c TSS, we performed a 5’RACE PCR on unstimulated and TCR-stimulated naïve CD4⁺ Tconv cells. A strong RACE band of ∼1kb was detected exclusively in stimulated cells (**Figure 2E**), with the 5’ end mapped by Sanger sequencing to the position reported in the FANTOM5 annotation at Chr11:111,512,478 (**Figure 2D – blue bar**). This TSS has been previously linked to miR-34b/c transcription in other cellular contexts (47). To assess the ability of FOXP3 to positively regulate miR-34c transcription, we cloned the genomic region around the identified TSS - nucleotides chr11:111,511,926-111,512,877 (forward strand) (**Figure 2D – black bar**) - containing two predicted FOXP3 binding sites, into the pGL4.10[*luc2*] reporter vector. We then performed dual-luciferase assays, with or without co-expression of recombinant FLAG-FOXP3. The reporter construct showed significant FOXP3-dependent activation, with luciferase levels increasing 65-fold when compared to the control assay performed without overexpression of the TF (**Figure 2F and 2G**). These findings indicate that FOXP3 can directly bind to this genomic region and activate a downstream promoter.

Together, these results provide strong evidence that the transcription of the miR-34b/c gene *locus* can be directly regulated by FOXP3, providing a justification for our observations that only thymic/natural Treg and naïve-derived cells expressing FOXP3⁺⁺ accumulate high levels of miR-34c following TCR stimulation (**Figure 1C and 2C**).

### miR-34c regulates genes associated with T cell activation and differentiation

To investigate the functional role of miR-34c, we first blocked its accumulation following TCR stimulation in purified human naïve CD4^+^ Tconv. For this purpose, we adapted our previously established method using cholesterol-conjugated antagomiRs (16) and set-up a robust protocol to prevent miR-34c upregulation following *in vitro* TCR stimulation. miR-34c levels significantly increased following TCR stimulation in control assays with a scramble antagomiR (AScr), while qPCR quantification revealed almost undetectable levels in cells stimulated in the presence of the miR-34c-specific antagomiR (A34c), up to 140 hours post-stimulation **(Figure 3A and 3B)**. The antagomir-dependent inhibition was found to be highly consistent, despite the inherent heterogeneity in TCR-driven activation responses across donors, which impacted miR-34c expression levels **(Figure 3A)**, as previously described (16).

**Figure 3.**
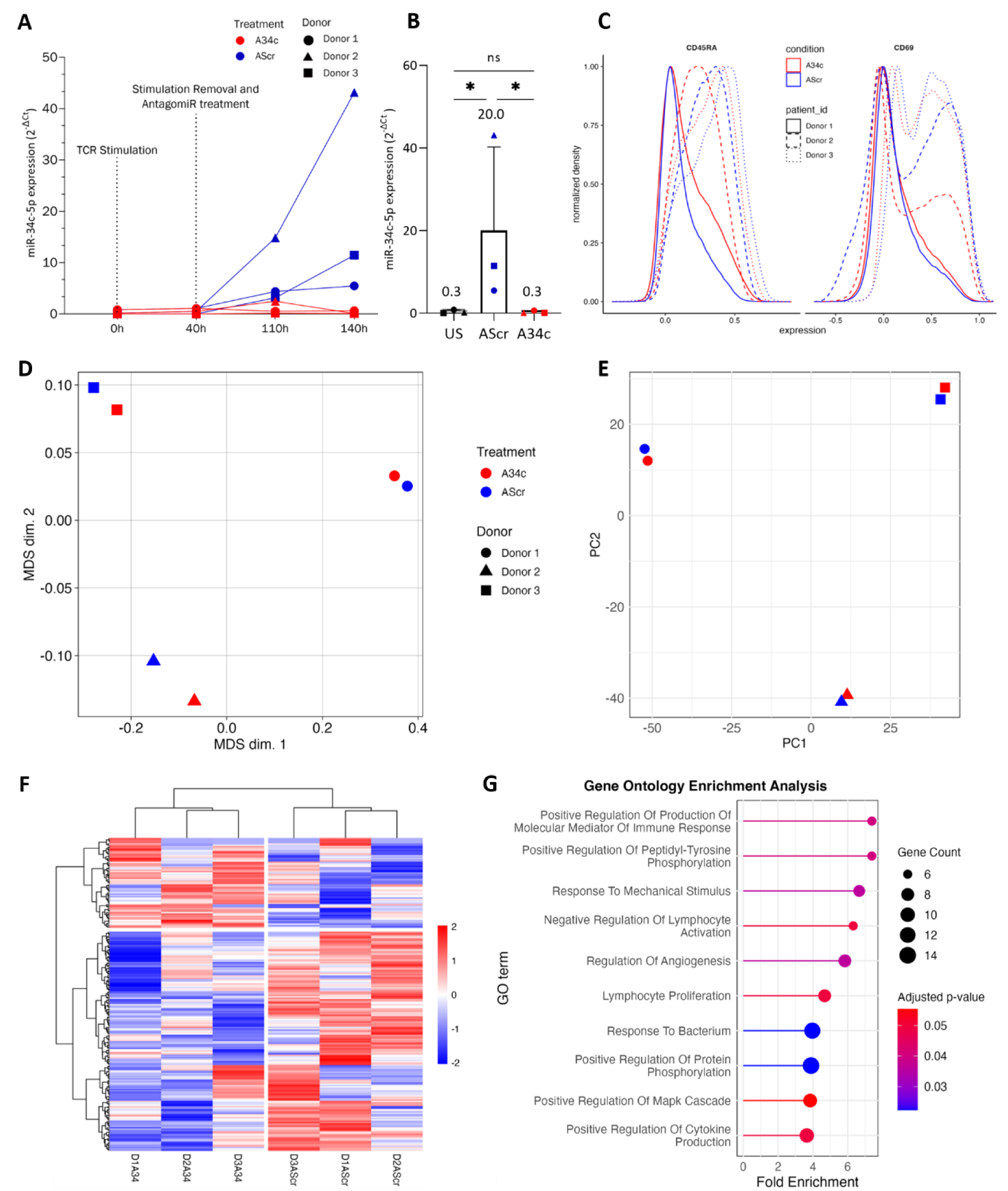
miR-34c regulates genes associated with T cell activation and differentiation. **(A)** Time course of miR-34c-5p expression following TCR stimulation of naïve CD4⁺ Tconv cells. Cells were stimulated with anti-CD3/CD28 and 10 U/mL IL-2 for 40 h, followed by stimulation removal and treatment with either a scrambled antagomiR (AScr - blue) or miR-34c-specific antagomiR (A34c - red) up to 140 h. Data is expressed as mean 2^−ΔCt^ to miR-191-5p ± SD. N = 3 independent donors. **(B)** miR-34c expression levels in unstimulated (US) and stimulated cells treated with AScr or A34c at 140 h post-stimulation. Data is expressed as mean 2^−ΔCt^ to miR-191-5p ± SD; * - p-value < 0.05 (one-tailed paired t-test). **(C)** Flow cytometry analysis of CD45RA and CD69 expression in AScr- and A34c-treated cells 140 h post-stimulation. Normalized density plots show expression profiles for three independent donors. **(D)** MDS of flow cytometry data **(E)** and PCA of RNA-seq data from A34c- and AScr-treated cells. Each point represents an individual sample, color-coded by treatment and shaped by donor. **(F)** Heatmap with hierarchical clustering of the DEGs identified in A34c- vs. AScr-treated cells (p-value ≤ 0.05, |log2foldchange| ≥ 0.6). Color scale represents mean log2-transformed expression level across samples. **(G)** Gene Ontology (GO) enrichment analysis of the DEGs in A34c- vs. AScr-treated cells (p-value ≤ 0.05, |log2foldchange| ≥ 0.6). Top enriched GO terms related to immune response regulation are shown, with dot size representing the number of genes associated with each term, the bar length showing the fold change, and color indicating adjusted p-values (FDR was adjusted for using the Benjamini-Hochberg).

The functional consequences of miR-34c knockdown were assessed by the analysis of cellular phenotypes by spectral flow cytometry coupled with transcriptome profiling using Illumina bulk mRNA sequencing. The flow cytometry analysis did not reveal any statistically significant differences on the expression of the main markers of cell activation/differentiation (CD3, CD4, CD45RA, CD45RO, CD25, CXCR3, CD69, ICOS, CXCR3, CD95, CCR7, CD31 and Live Dead – data not shown). Analysis of CD45RA and CD69 intensity distributions highlighted the large variability in expression across donors, which might obscure the antagomir-specific effect **(Figure 3C)**. In agreement with this, multidimensional scaling (MDS) analysis of the flow cytometry activation marker panel revealed that samples grouped primarily by donor rather than by treatment **(Figure 3D)**, suggesting that inter-donor variability outweighed any global effects of miR-34c knockdown on T cell activation.

We further characterized the transcriptome profile of each sample, sequenced to an average depth of 80 million paired-end reads (ranging from 65 to 120M reads), allowing us to detect ∼19,000 expressed genes (**Supplementary file S1**). Exploratory analysis of the mRNA-seq data revealed a profile closely mirroring the MDS analysis of the flow cytometry activation panel, with the first two principal components explaining ∼95% of the total variance (**Figure 3E**). This implies that donor-specific effects strongly dominate the expression profiles, obscuring the more subtle effects of miR-34c inhibition. To address these limitations, we performed differential expression analysis using the limma-voom pipeline with a paired design, applying a relaxed threshold of |log₂ fold-change| ≥ 0.6 and nominal p-value ≤ 0.05 to identify candidate genes potentially modulated by miR-34c. This approach identified 234 genes that responded to miR-34c inhibition, of which 74 were upregulated, and 160 were downregulated (**Supplementary Figure 3** and **Supplementary file S2**). Unsupervised hierarchical clustering analysis of this gene set revealed a clear separation of samples based on treatment with miR-34c-antagomiR versus scrambled-antagomiR. Despite pronounced inter-donor variability and some gene-level inconsistencies, the clustering grouped both genes and conditions into two main clusters corresponding to those upregulated or downregulated in response to antagomiR treatment (**Figure 3F**). To gain further insights into the functional role of miR-34c in CD4⁺ T cells, we performed a Gene Ontology (GO) enrichment analysis using the 234 genes that responded to the inhibition of this miRNA. Several immune-related functional signatures were identified (**Figure 3G**), suggesting that this miRNA influences key T cell biological processes. These included processes related to the production of immune response mediators, lymphocyte activation, proliferation and regulation of cytokine production. Despite the inter-donor variability and the limited sensitivity of transcriptome profiling for detecting strong miRNA-dependent signatures, these results suggest that miR-34c induction likely plays a relevant role in T cell activation following TCR stimulation.

### miR-34c limits iTreg differentiation by restraining FOXP3 expression

Given our observation that high miR-34c expression is restricted to cells with high levels of FOXP3, it is possible that its effects would become more pronounced in an experimental setting favoring the differentiation of FOXP3^+^ cells. In fact, the restricted expression of miR-34c to only a few cell subsets in our experimental system likely reduced our ability to detect clear phenotypic effects following its inhibition during TCR stimulation. To refine our analysis, we employed an optimized stimulation protocol designed to promote Treg cell differentiation. We stimulated naïve CD4^+^ T cells, previously treated with miR-34c or scrambled antagomiRs, in the presence of TGF-β and IL-2 for 120 hours. These conditions are well-established to induce FOXP3 expression and iTreg development (12). Additionally, we expanded our spectral flow cytometry panel to better evaluate the functional impact of miR-34c modulation under these conditions (CD3, CD4, CD45RA, CD45RO, CD25, FOXP3, CXCR3, CD69, ICOS, CXCR3, CCR7, CD127, Helios, CD39, CCR6, OX40, PD-1, PD-L1, CTLA-4, ki-67, CXCR5 and Live Dead).

miR-34c knockdown significantly increased the frequency of FOXP3⁺CD25⁺ cells, as shown in representative flow cytometry plots **(Figure 4A)** and quantified across three independent donors **(Figure 4B)**. These results suggest that miR-34c negatively regulates iTreg differentiation. Importantly, miR-34c knockdown did not affect proliferation, as assessed by Ki-67 staining **(Supplementary Figure 4)**, in contrast to previous reports in other cellular systems (48). In addition to altering the frequency of FOXP3⁺ cells, miR-34c knockdown also enhanced FOXP3 protein levels. Histogram of FOXP3 fluorescence intensity show a clear shift in miR-34c-antagomiR-treated cells compared to the control **(Figure 4C)**, which was consistent across donors, as shown by the quantification of FOXP3 mean fluorescence intensity (MFI) in three independent donors **(Figure 4D)**. This increase in protein expression was also modestly reflected at the transcript level, with *FOXP3* mRNA levels elevated by 13% and 27% in two of three donors in our RNA-seq dataset **(Supplementary Figure 5)**.

**Figure 4.**
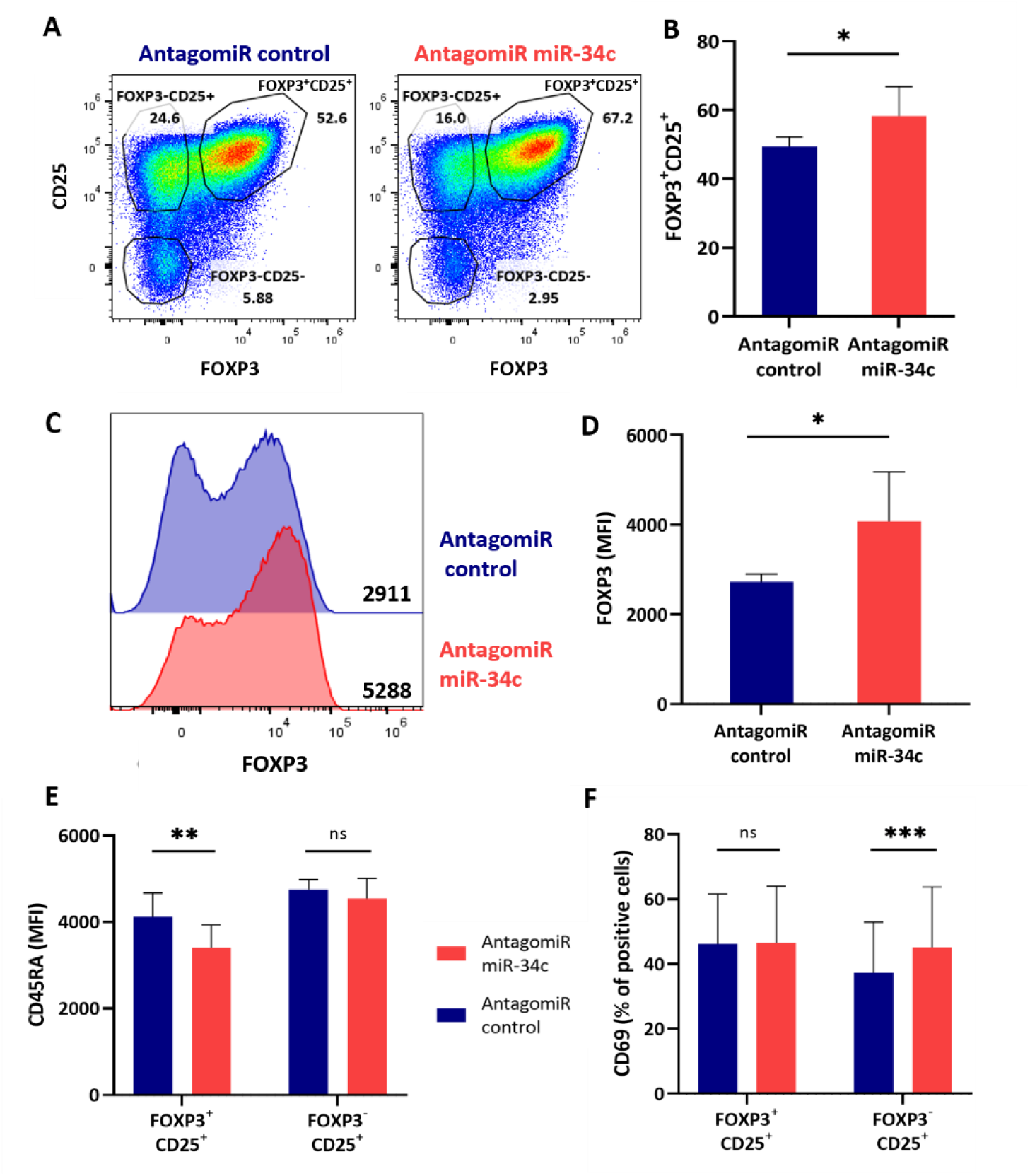
miR-34c regulates T cell activation and differentiation in a subset-specific manner. **(A)** Representative flow cytometry plot of CD25 (IL2RA) and FOXP3 in A34c- and AScr-treated naïve CD4^+^ T cells after 120 h of TCR stimulation conditions to promote iTreg differentiation. **(B)** Quantification of FOXP3⁺CD25⁺ cell population in three biological replicate assays. Data is represented as mean ± SD; * - p-value < 0.05 (one-tailed paired t-test). **(C)** Histogram of a representative MFI distribution of FOXP3 in A34c- and AScr-treated naïve CD4^+^ T cells in the samples from panel 4A, and **(D)** the correspondent quantification plot of the MFI of FOXP3 in three independent experiments. Data is represented as mean ± SD; * - p-value < 0.05 (two-tailed paired t-test). **(E)** CD45RA MFI within FOXP3⁺CD25⁺ and FOXP3⁻CD25⁺ populations following miR-34c knockdown. Data is represented as mean ± SD; ns – not significant; ** - p-value < 0.01 (two-tailed paired t-test). **(F)** Percentage of CD69⁺ cells within FOXP3⁺CD25⁺ and FOXP3⁻CD25⁺ populations after miR-34c knockdown. Data is represented as mean ± SD; ns – not significant; *** - p-value < 0.001 (two-tailed paired t-test).

We next investigated the expression of activation and differentiation markers within FOXP3⁺CD25⁺ and FOXP3⁻CD25⁺ subsets. CD45RA expression, a marker of naïve or undifferentiated T cells, was significantly reduced within the FOXP3⁺CD25⁺ subset following miR-34c knockdown, while it remained unchanged in FOXP3⁻CD25⁺ cells **(Figure 4E)**. This suggests that miR-34c may contribute to maintaining cells in a more naïve state and that its inhibition facilitates iTreg differentiation. In contrast, CD69 expression showed a distinct pattern. While no significant change was observed within the FOXP3⁺CD25⁺ population, a significant increase in CD69⁺ cells was detected in the FOXP3⁻CD25⁺ subset after miR-34c knockdown **(Figure 4F)**. These findings support a model in which miR-34c exerts context-dependent regulatory functions in distinct T cell subsets, potentially targeting different transcripts depending on the differentiation state. Together, these results indicate that miR-34c weakens iTreg differentiation by repressing FOXP3 expression and maintaining a more naïve or less activated phenotype. Its inhibition enhances both the frequency and expression level of FOXP3⁺ iTregs while differentially modulating early activation markers in a subset-specific manner.

### miR-34c inhibits FOXP3 expression by targeting its 3’UTR

Given that miR-34c knockdown led to an increase in FOXP3 protein levels, we then investigated whether miR-34c directly regulates *FOXP3* at the post-transcriptional level. We used the RNA22 miRNA target prediction algorithm (49) to scan the *FOXP3* 3’ UTR for putative miR-34c binding sites. RNA22 identified four putative miR-34c miRNA response elements (MREs), two of which separated by a stretch of ∼50nt, thereby representing a strong candidate inhibitory element (50) **(Figure 5A)**. To experimentally validate *FOXP3* as a *bona-fide* direct target of miR-34c, we cloned a portion of the *FOXP3* 3′UTR containing the four predicted MREs into a luciferase reporter vector (*FOXP3* 3′UTR) and performed dual-luciferase reporter assays in HEK-293 cells. A second reporter vector with point mutations predicted by RNA22 to abrogate the interaction of miR-34c with these sites (*FOXP3* 3′UTR-mut) was generated as a control. Co-transfection of miR-34c with the *FOXP3* 3′UTR construct resulted in a significant reduction in luciferase activity, while mutation of the binding sites abolished this effect **(Figure 5B)**. These results demonstrate that miR-34c directly regulates FOXP3 expression levels by interacting with specific MREs in the 3’UTR of the mRNA.

**Figure 5.**
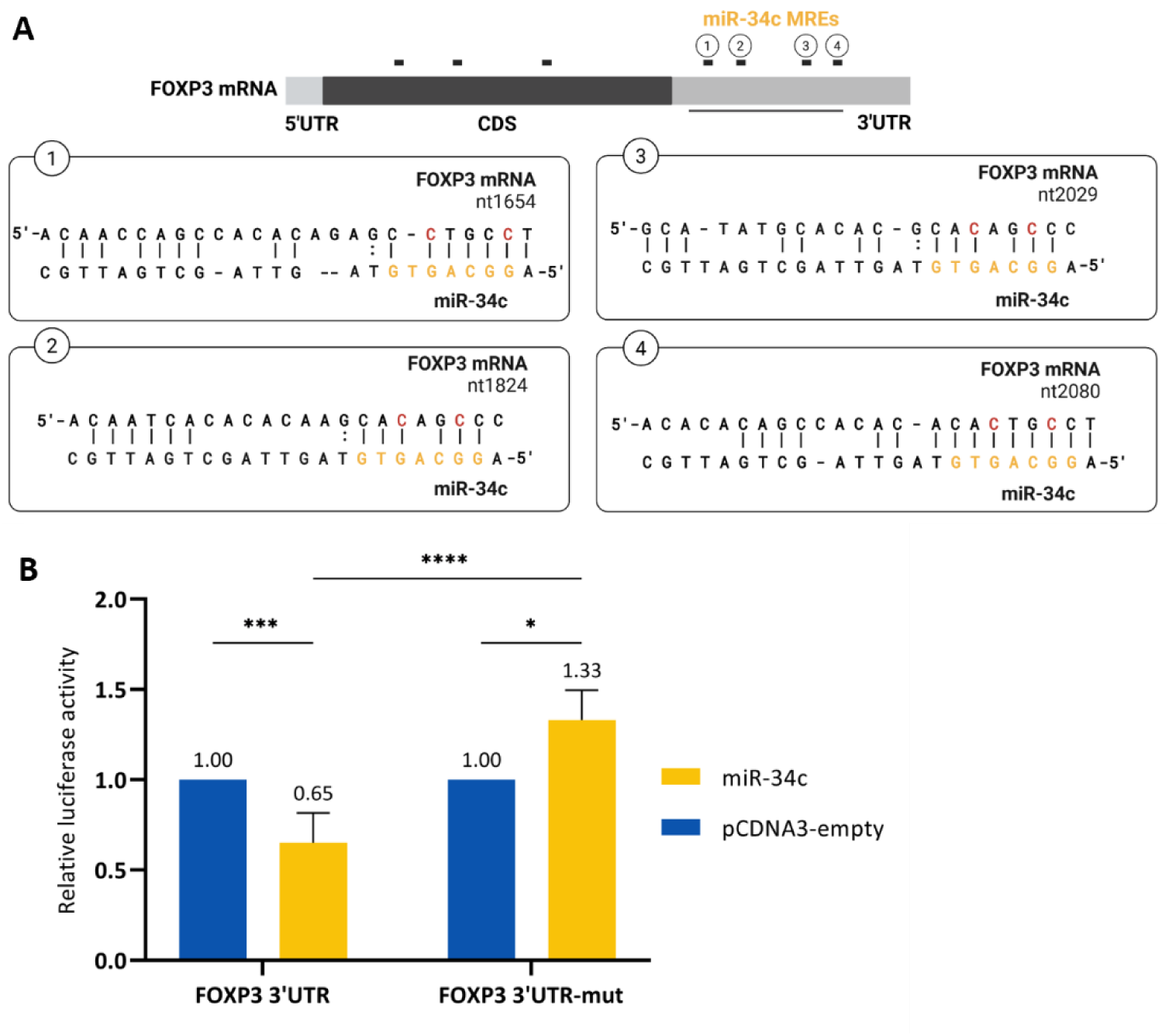
miR-34c inhibits *FOXP3* expression by targeting its 3’UTR. **(A)** Predicted MREs for miR-34c in the *FOXP3* mRNA (black bars) and predicted miRNA-MRE hybridization for the four MREs located in the 3’UTR. Nucleotides in yellow represent miR-34c’s seed sequence while nucleotides in red illustrate the positions mutated to prevent miR-34c from binding to the *FOXP3* 3’UTR. *FOXP3* 3’UTR-mut was built by replacing the red-highlighted cytosines for adenines **(B)** Dual luciferase reporter assay to analyze the interaction of miR-34c with the *FOXP3* 3’UTR. The 3’ UTR of *FOXP3* and the mutated version (*FOXP3* 3’UTR-mut) were cloned in a dual-luciferase reporter vector downstream of the firefly luciferase coding sequence and these were transfected into HEK-293 cells together with either pcDNA3-miR-34c or pcDNA3-empty control. Firefly luciferase luminescence was normalized to that of the renilla luciferase. N=3 to 6 independent experiments. Data is represented as mean ± SD; * - p-value < 0.05; *** - p-value < 0.001; **** - p-value < 0.0001 (two-way ANOVA).

Taken together, our data provides a model whereby following TCR activation, miR-34c expression is strongly induced in FOXP3^+^ cells and establishes a negative feedback loop by targeting *FOXP3* mRNA, acting as a negative modulator of iTreg differentiation.

## Discussion

In this work we identified miR-34c as a novel regulator of iTreg differentiation, whose expression was specifically induced as part of the response of naïve CD4^+^ T cells to TCR stimulation. Our results showed that miR-34c expression was sustained in FOXP3⁺ cells, and that this transcription factor acted both as an activator and a target of miR-34c. We propose a model whereby miR-34c is involved in a negative regulatory feedback loop that is relevant for the establishment of the iTreg cell population.

miR-34c is part of a conserved family of six homologous miRNAs (miR-34a, miR-34b/c, and miR-449a/b/c) encoded at three distinct genomic loci (51). These miRNAs have been reported to act redundantly on multiple processes including the control of motile ciliogenesis, spermatogenesis, cellular reprogramming, and brain development, with no phenotypes reported in single-gene knockout mice (52). However, in our previous work, we found that miR-34c-5p is the only member of this family induced in response to TCR stimulation in naïve CD4^+^T cells (16). This implies a specific, non-redundant role for this miRNA in immune system function, which we addressed in the present study.

Our analysis of the *ex vivo* miR-34c expression in sorted T cell populations – naïve and memory Tregs, naïve Tconv, Th1, Th2, and Th17 – showed that *ex vivo* miR-34c was exclusively detected in memory Treg cells. This observation is supported by publicly available data from the Database of Immune Cells (DICE) (44), in which miR-34c primary transcript is detected exclusively in memory Tregs, although previous studies addressing the Treg-specific miRNA signature did not report its expression (8,53). Interestingly, upon TCR stimulation miR-34c was strongly upregulated in both Treg and Tconv naïve CD4⁺ cells, while its expression remained unchanged in memory Tregs and was undetectable in conventional memory subsets. This contrasts with the broad induction of other activation-induced miRNAs like miR-155, revealing that miR-34c expression is not a general feature of T cell activation, and may interplay with *FOXP3* to modulate the conventional versus regulatory differentiation upon TCR stimulation of naïve CD4^+^ T cells.

We hypothesize that the observed distinct behaviour of naïve and memory CD4⁺ T cells may reflect underlying differences in chromatin regulation and responsiveness to TCR stimulation. TCR engagement induces substantial epigenetic remodeling in both naïve and memory T cells, but the dynamics and extent of these changes differ (reviewed in (54,55)). In naïve T cells, activation involves the recruitment of chromatin remodeling complexes, progressive demethylation of promoter regions, increased histone acetylation, and deposition of activating histone marks such as H3K4me3, changes that collectively enable transcription factor binding and transcriptional activation (56). In contrast, memory T cells exhibit pre-established chromatin accessibility and enrichment of active histone modifications at many effector loci, enabling a faster and more robust transcriptional response upon restimulation (56–58). A recent genome-wide ATAC-seq study comparing thymic and peripheral Treg and Tconv cells reported no differences in chromatin accessibility at the miR-34c regulatory region, which was consistently in an open state (59). However, ENCODE ChIP-seq data revealed markedly higher levels of H3K4me3 at this locus in Treg cells (22), suggesting additional layers of regulation through histone modifications (see **Supplementary figure 6**). Although a detailed investigation of naïve and memory T cell responses lies beyond the scope of this study, these observations support the idea that differences in epigenetic priming and remodeling capacity may underlie the differential induction of miR-34c in response to TCR stimulation.

Consistent with our findings that miR-34c is expressed in Treg cells, a previous study has reported a reduction in the circulating levels of miR-34c in two independent models of Treg depletion (FOXP3-DTR mice and anti-CD25 antibody-treated mice), as well as in patients with autoimmune diseases including rheumatoid arthritis, systemic lupus erythematosus, Sjögren’s syndrome, and ulcerative colitis (60). These findings provide compelling evidence that miR-34c levels in circulation reflect Treg abundance or functional integrity *in vivo*, pointing to its potential as a biomarker for immune homeostasis.

We showed that FOXP3 can activate the transcription of miR-34c and was simultaneously a direct target of miR-34c-mediated repression via 3′UTR interaction. This regulatory architecture suggests a negative feedback loop in which miR-34c modulates the abundance of FOXP3 - the central transcription factor controlling Treg differentiation, function, and identity. A related example is miR-31, which was first identified as a direct post-transcriptional regulator of *FOXP3* via its 3′UTR (8), and later shown to regulate Treg differentiation, independently of *FOXP3*, through *GPRC5A*, a molecule associated with retinoic acid signaling (7). Interestingly, FOXP3 also binds directly to the promoter of miR-31, suppressing its expression following TCR activation, and thereby forming a reciprocal negative feedback loop. In contrast, miR-34c and FOXP3 appear to establish a unidirectional feedback mechanism, in which FOXP3 promotes miR-34c expression and is in turn repressed by it. These observations underscore the existence of multiple microRNA-mediated feedback circuits converging on FOXP3 to fine-tune Treg differentiation and function. Coupled to the dynamic nature of immune cell populations, this complex regulatory network makes the investigation of specific miRNA functions particularly difficult. The use of antagomiRs to inhibit miR-34c expression in bulk Tconv stimulation experiments did not reveal strong global phenotypic effects. This likely reflects a combination of donor variability, the low abundance of FOXP3⁺ cells in bulk cultures, and the context-dependent action of miR-34c. By shifting to TGF-β–induced iTreg conditions, we were able to reveal a specific phenotype upon miR-34c knockdown, suggesting that the functional impact of this miRNA becomes evident in a permissive differentiation environment.

In addition to the feedback loop established with FOXP3, it is noteworthy that the transcriptional co-activator *PCAF*/*KAT2B*, a well-characterized miR-34c target, is also involved in the control of iTreg differentiation (16,61,62). Loss of PCAF impairs the development of iTreg cells both *in vitro* and *in vivo*. Specifically, PCAF−/− CD4^+^ T cells show reduced conversion to iTregs, with lower proportions of CD4^+^FOXP3^+^ cells and decreased *Foxp3* mRNA expression. When Tregs from PCAF−/− mice are activated via TCR stimulation, they show significantly lower proportions of CD4^+^FOXP3^+^ cells and increased apoptosis compared to controls (62). This phenotype is the opposite of what we observe upon knockdown of miR-34c, supporting a role for miR-34c in a regulatory network that limits iTreg differentiation via repression of positive regulators.

We detected miR-34c expression in FOXP3⁻CD25^+^ cells following TCR stimulation under iTreg-inducing conditions, albeit at lower levels than in FOXP3^+^ cells. In this setting, FOXP3^⁻^CD25^+^ cells may represent recently activated conventional cells, in line with their high levels of CD69 without changes in CD45RA expression. The up-regulation of miR-34c may play a role in limiting the *FOXP3* upregulation and the full-differentiation into iTregs, which have already lost the expression of CD45RA and down-regulated CD69.

Taken together, our data supports a model in which miR-34c acts as a context-dependent modulator of iTreg differentiation through both transcriptional and post-transcriptional regulation. Its selective induction in FOXP3⁺ cells and its ability to fine-tune FOXP3 expression suggest that miR-34c may function as a post-transcriptional rheostat controlling the threshold for Treg lineage commitment and stability, with potential implications for immune tolerance and autoimmunity.

## Acknowledgements

Authors thank IPST for providing the buffy coats and the Flow Cytometry Platform of Gulbenkian Institute for Molecular Medicine for their technical support in the sorting experiments. The authors acknowledge Mark Gibson for his role in establishing primary T cell manipulation and transfection methods in the Gama-Carvalho lab. The authors also thank current members of the Sousa Lab, Diana Santos, Alexandre Raposo, and Afonso Almeida; as well as former members Ana Godinho-Santos, Iris Caramalho, and André Gomes, for their valuable technical support and insightful discussions throughout this work. This work was funded by the following grants from Fundação para a Ciência e a Tecnologia (FCT), Portugal, through PTDC/BIA-CEL/29257/2017 and by UID/04046/2025 - Instituto de Biosistemas & Ciências Integrativas Centre grant. CN-E is funded by the fellowship 2020.05340.BD from FCT, Portugal.

## Author Contributions

CN-E, MG-C and AES conceived and designed the study. CN-E and RG-L performed experiments with primary cells. CN-E analyzed data. GP-O and TFA analyzed RNA-seq data with assistance from MG-C. TFA and CF-F performed FOXP3 luciferase assays. MP-P cloned miR-34c regulatory region in the luciferase reporter vector and established the experimental conditions for 5’RACE PCR. MPM assisted CN-E in the generation of samples for RNA-seq and the establishment of flow cytometry panels. CN-E wrote the article. MG-C contributed to writing the Discussion and prepared the figures related to RNA-seq analysis. CN-E prepared all other figures. MG-C and AES secured funding and supervised the work. All authors reviewed and approved the final manuscript.

## Figure legends

**Supplementary figure 1.**
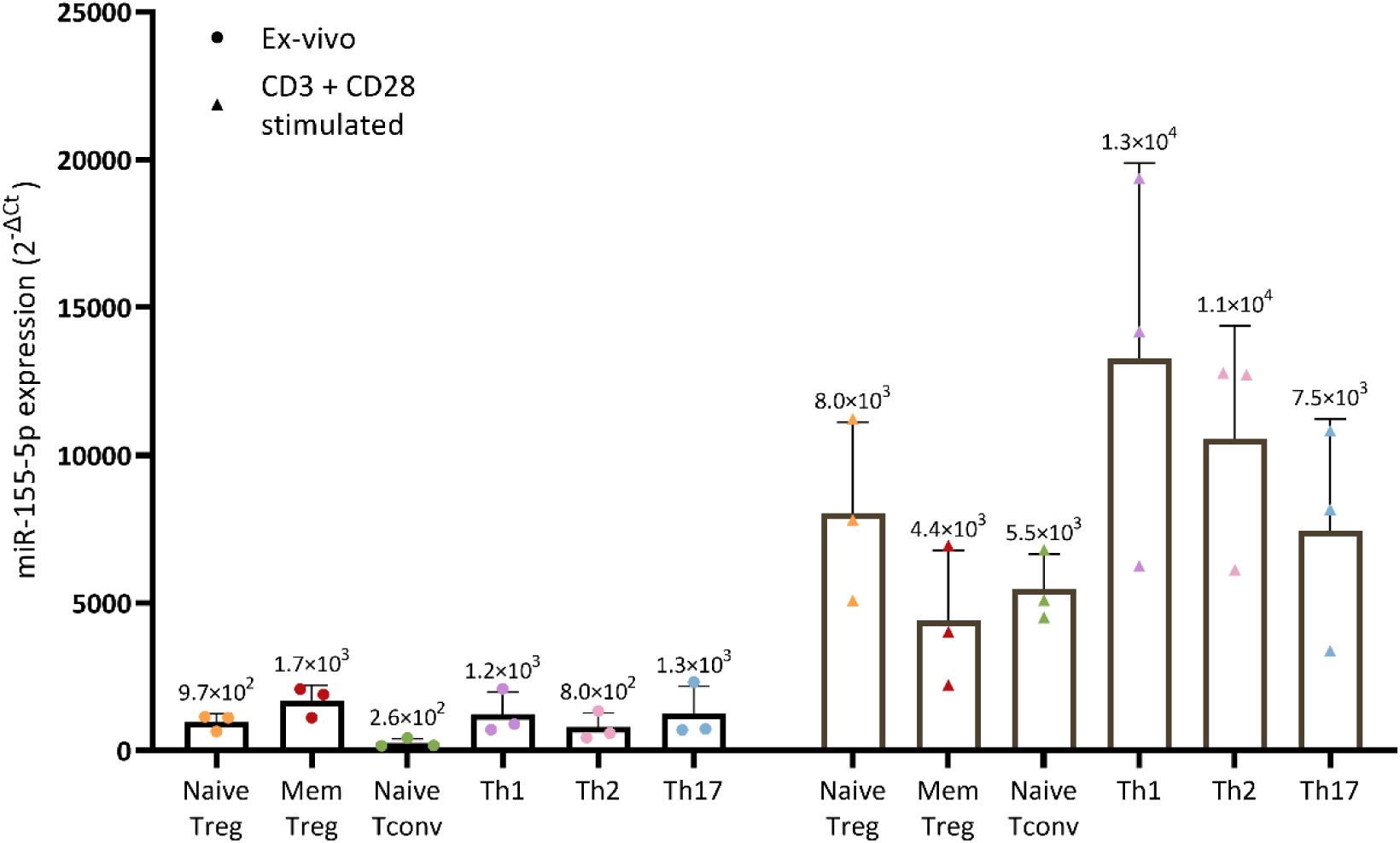
miR-155-5p expression levels measured by RT-qPCR in *ex vivo* (circle shape) or after *in vitro* CD3/CD28 stimulation (triangle shape) of various sorted CD4⁺ T cell subsets. Subsets include naïve regulatory T cells (Naive Treg), memory regulatory T cells (Mem Treg), naïve conventional T cells (Naive Tconv), and effector memory Th subsets (Th1, Th2, Th17). Data is expressed as mean 2^−ΔCt^ to miR-191-5p ± SD. N = 3 independent donors.

**Supplementary figure 2.**
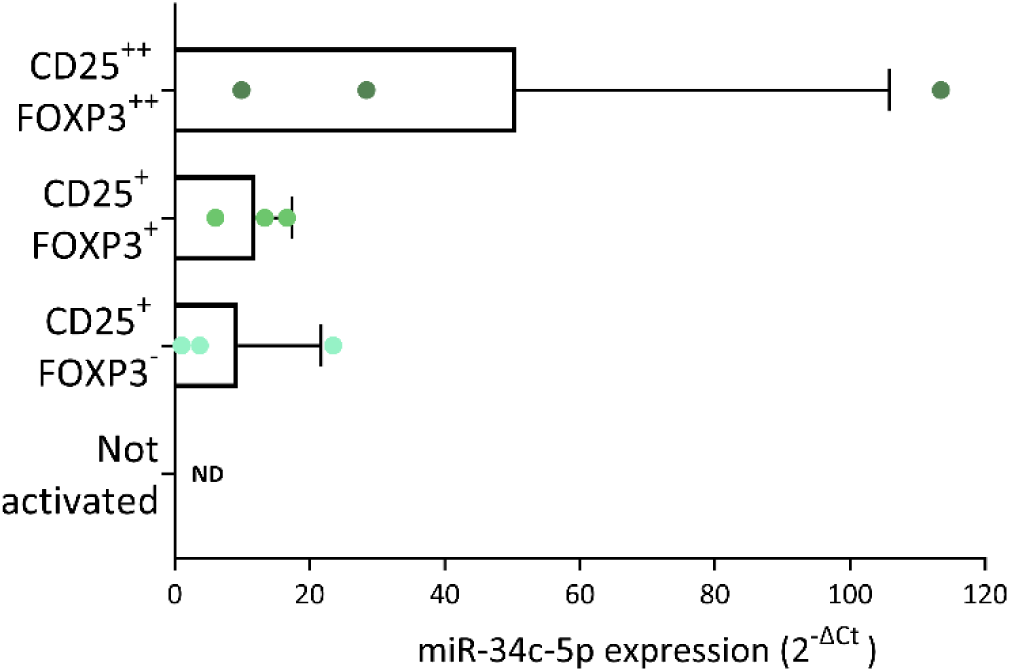
miR-34c expression levels measured by RT-qPCR in sorted CD4⁺ T cell subsets with varying CD25 and foxp3 expression levels. Naïve T conv cells were stimulated with CD3/CD28 Dynabeads + IL-2 + TGF-β for 120 h before sorting and miR-34c quantification. Subsets include CD25|FOXP3 high (++), CD25|FOXP3 medium (+) CD25 medium|FOXP3 negative (-) and CD25|FOXP3 negative cells. Data is expressed as mean 2^−ΔCt^ to miR-191-5p ± SD. N = 3 independent donors.

**Supplementary figure 3.**
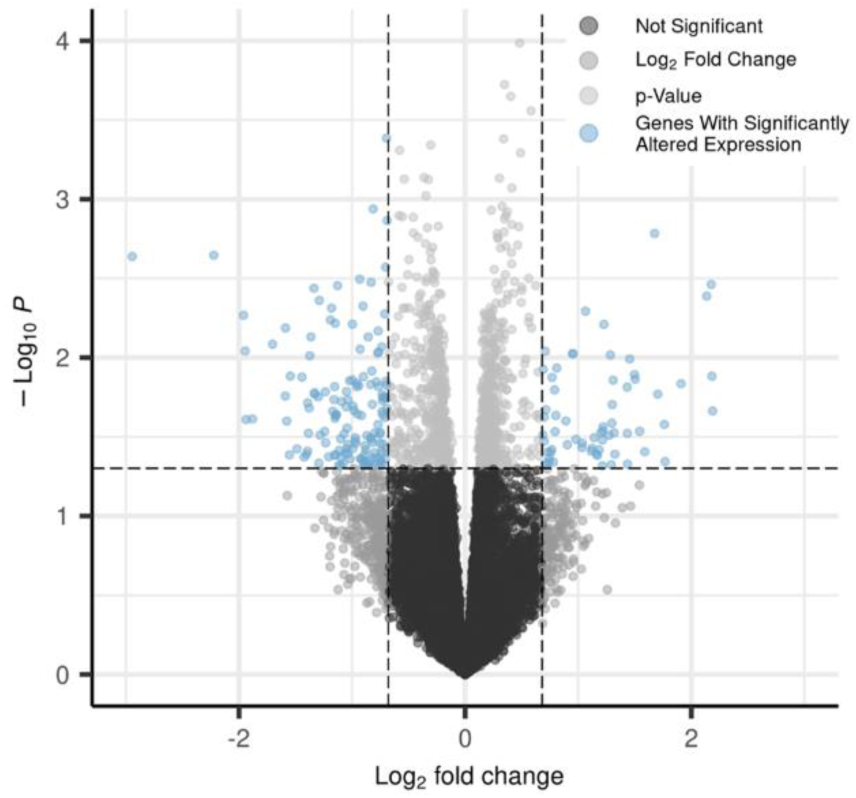
Volcano plot of the DEGs upon miR-34c knockdown. The volcano plot displays the results of the paired DGE analysis between the scrambled-antagomiR (control) and the miR-34c-antagomir-treated samples. The x-axis represents the log2 fold change of normalized gene expression, with positive values indicating upregulation and negative values indicating downregulation upon miR-34c knockdown. The y-axis shows the negative log10 of the p-value, indicating statistical significance.

**Supplementary figure 4.**
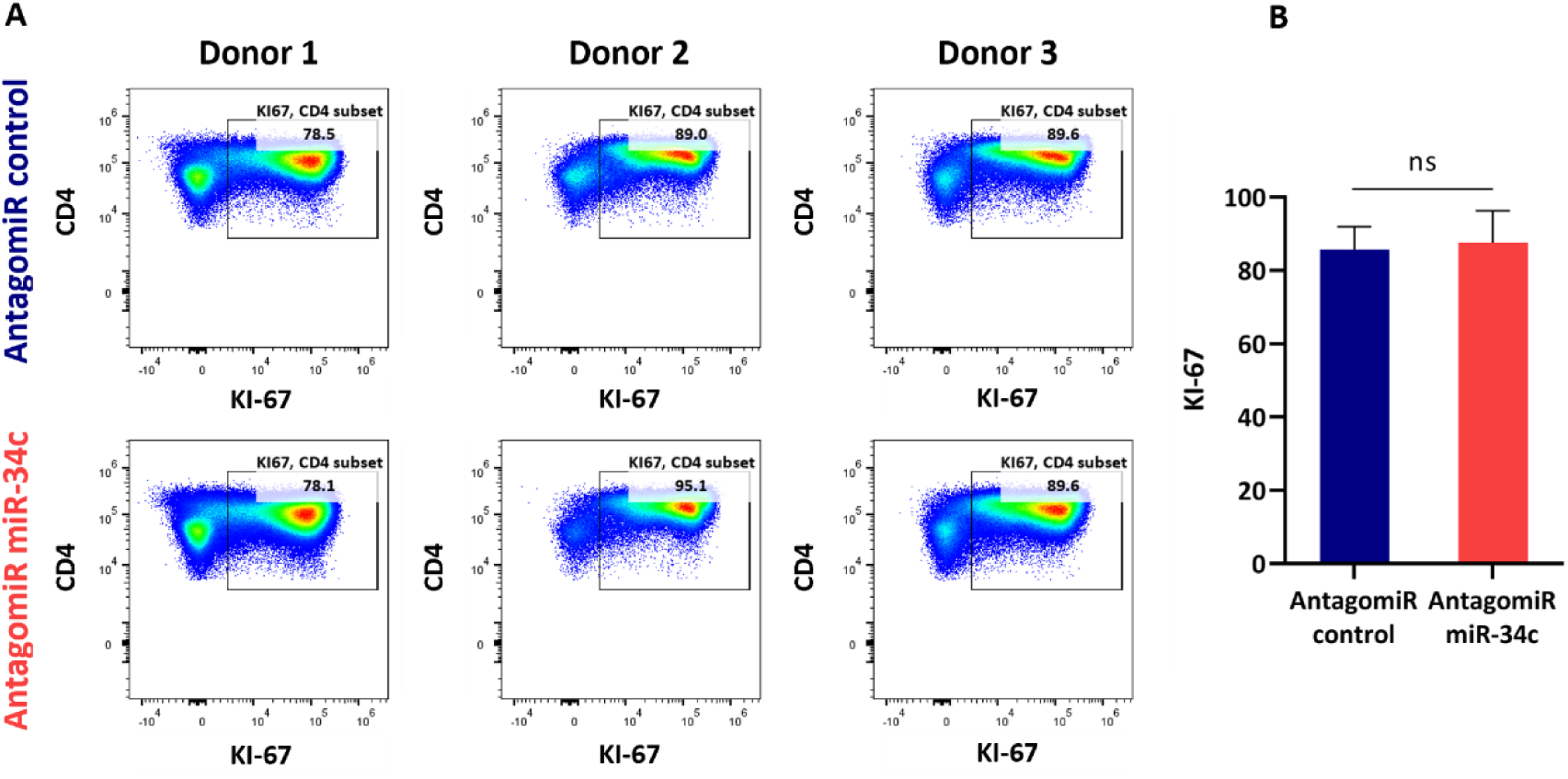
miR-34c does not impact the proliferation of CD4^+^ T cells **(A)** Flow cytometry plots showing the percentage of Ki-67^+^ cells within live CD3⁺CD4⁺. Naïve Tconv cells were treated with either scrambled AntagomiR (top row) or miR-34c AntagomiR (bottom row) followed by CD3/CD28 Dynabeads + IL-2 + TGF-β. **(B)** Bar chart showing the percentage of Ki-67⁺ CD4⁺ T cells from three biological replicates per group. Data is represented as mean ± SD; ns – not significant (two-tailed paired t-test).

**Supplementary figure 5.**
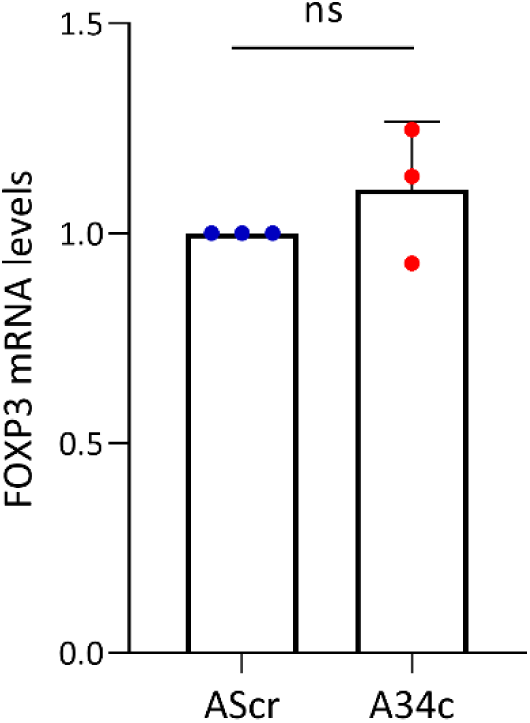
*FOXP3* expression levels from the RNA-seq experiment in naïve CD4⁺ Tconv cells treated with AScr or A34c. Data is normalized to AScr. Data is represented as mean ± SD; ns – not significant (two-tailed paired t-test).

**Supplementary figure 6.**
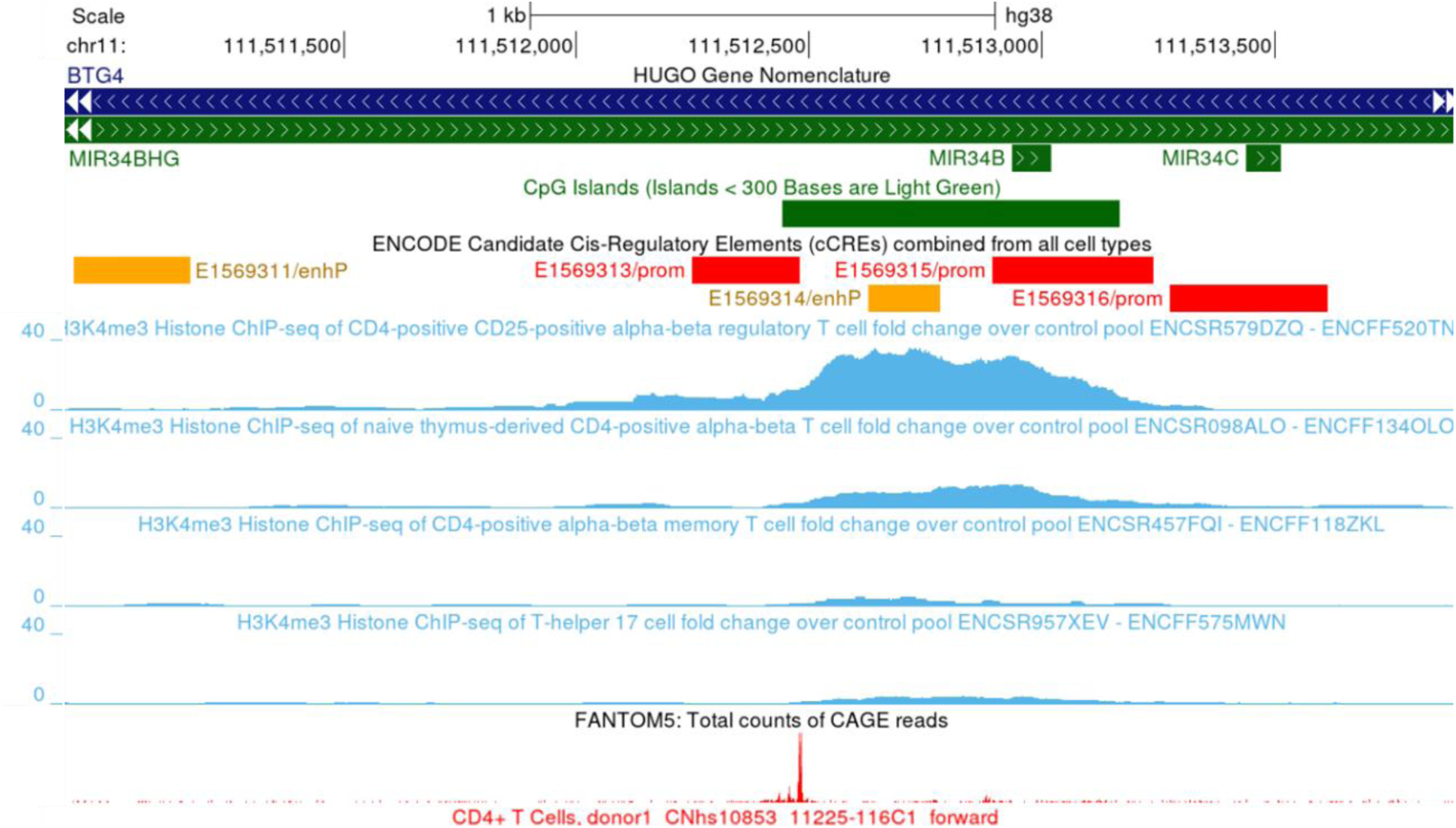
The *MIR34C locus* in the GRCh38 assembly, annotated on the UCSC Genome Browser with ENCODE, and FANTOM5 data. Displayed are annotated promoter and enhancer regions (red and yellow bars), CpG island from ENCODE (green bar), and a TSS from FANTOM5 CD4^+^ T cell CAGE datasets. ENCODE Histone H3K4me3 ChIP-seq signal (light blue peaks) across the *MIR34C locus* in several human CD4⁺ T cell subsets are also displayed. ChIP-seq tracks show H3K4me3 fold enrichment (pooled fold change over input) in CD25⁺ Tregs, naïve thymus-derived CD4⁺ T cells, memory CD4⁺ T cells, and Th17 cells.

