## Supplementary material for "miR-34c-5p is a novel regulator of T cell differentiation that targets FOXP3": Table 1

| Target | Clone | Fluorophore | Source | Catalog number | Dilution | Panel |
|---|---|---|---|---|---|---|
| CD3 | SK7 | APC-Fire 810 | Biolegend | 344858 | 1:150 | Panel 1, 2, 3, 4 |
| CD4 | RPA-T4 | PE-Dazzle | Biolegend | 300548 | 1:200 | Panel 1, 2, 4 |
| CD45RA | HI100 | BV650 | Biolegend | 304136 | 1:100 | Panel 1, 2, 4 |
| CD45RO | UCHL1 | BV570 | Biolegend | 304226 | 1:100 | Panel 1, 2, 3, 4 |
| CCR7 | G043H7 | BV711 | Biolegend | 353228 | 1:30 | Panel 1, 2, 4 |
| CD25 | BC96 | PE-Cy5 | Biolegend | 302608 | 1:150 | Panel 1, 2, 3, 4 |
| CD69 | FN50 | Spark-NIR 685 | Biolegend | 310958 | 1:200 | Panel 1, 2, 3, 4 |
| CD127 | A019D5 | Alexa Fluor 700 | Biolegend | 351344 | 1:50 | Panel 1, 2, 4 |
| CXCR3 | G025H7 | PE-Cy7 | Biolegend | 353720 | 1:120 | Panel 1, 2, 3, 4 |
| CXCR5 | 51505 | APC | R&D Systems | FAB190A-100 | 1:20 | Panel 2 and 4 |
| CCR4 | 1G1 | BV750 | BD | 746980 | 1:100 | Panel 1, 2, 4 |
| ICOS | C398.4A | BV510 | Biolegend | 313525 | 1:200 | Panel 2 and 4 |
| PD-1 | EH12.2H7 | BV605 | Biolegend | 329924 | 1:150 | Panel 2 and 4 |
| PD-L1 | 29E.2A3 | BV785 | Biolegend | 329736 | 1:100 | Panel 2 and 4 |
| Helios | 22F6 | Alexa Fluor 488 | Biolegend | 137223 | 1:50 | Panel 2 and 4 |
| CD4 | RPA-T4 | V500 | BD | 560768 | 1:60 | Panel 3 |
| CD31 | WM59 | BV785 | Biolegend | 303148 | 1:75 | Panel 3 |
| CCR7 | 150503 | FITC | R&D Systems | FAB197F-100 | 1:15 | Panel 3 |
| CD45RA | HI100 | PerCP-Cy5.5 | Invitrogen | 45-0458-42 | 1:75 | Panel 3 |
| CD95 | DX2 | PE | Invitrogen | 12-0959-42 | 1:100 | Panel 3 |
| ICOS | C398.4A | Alexa Fluor 700 | Biolegend | 313528 | 1:150 | Panel 3 |
| OX40 | Ber-ACT35 (ACT35) | BV421 | Biolegend | 350014 | 1:150 | Panel 2 and 4 |
| CTLA-4 | BNI3 | PE | BD | 555853 | 1:100 | Panel 2 and 4 |
| CD39 | eBioA1 | PerCP-eF710 | Invitrogen | 46-0399-42 | 1:150 | Panel 2 and 4 |
| Foxp3 | PCH101 | eF450 | eBioscience | 48-4776-42 | 1:50 | Panel 2 and 4 |
| CCR6 | 11A9 | BV480 | BD | 566130 | 1:50 | Panel 1, 2, 4 |
| Ki-67 | B56 | PerCP-Cy5.5 | BD | 561284 | 1:60 | Panel 2 and 4 |
| Live-Dead | N/A | NIR | Invitrogen | L34975 | 1:500 | Panel 1, 2, 3, 4 |
