## Supplementary material for "miR-34c-5p is a novel regulator of T cell differentiation that targets FOXP3": Table 2

| Name | Sequence (5' – 3') | Application |
|---|---|---|
| miR-34c prom_Fw | AGAGGACCACGATTTGATG | Amplify miR-34c regulatory region from genomic DNA |
| miR-34c prom_Rv | CCTCGGACCCCATTTCAC | Amplify miR-34c regulatory region from genomic DNA |
| Prom_XhoI_Fw | TAATACTCGAGAGAGGACCACGATTTGATG | Create sticky ends to clone miR-34c regulatory region into pGL4.10 vector |
| Prom_HindIII_Rv | TTACTAAGCTTCCTCGGACCCCATTTCAC | Create sticky ends to clone miR-34c regulatory region into pGL4.10 vector |
| FOXP3 UTR_Fw | TCAGTGTCTAGAAAGGAGGATGGACGAACAGG | Amplify FOXP3 3'UTR |
| FOXP3 UTR_Rv | CAGTCACTCGAGGGGGAGACACGGGGTATTTT | Amplify FOXP3 3'UTR |
| 5'RACE_miR-34c | ACCTGGCCGTGTGGTTAGTGATTGGT | Gene specific primer |
| Seq_RACE_miR-34c | TATTAGCAATCAGCTAACTACACTGCCTAG | RACE product amplification and sequencing |
| 2_Seq_RACE_miR-34c | CATTTCACCGGGGGACGCCCTACCATGGCT | Sequencing 5'UTR |
