## Supplementary file S1 for "miR-34c-5p is a novel regulator of T cell differentiation that targets FOXP3"

| Sample ID | Treatment | Donor | Library ID | Raw reads | Filtered reads | Single mapped reads | % of single mapped reads | Total number of genes detected (>0 counts) |
|---|---|---|---|---|---|---|---|---|
| D1A34 | miR-34c antagomiR | #1 | D1A34_12739AAD_CGTTGGTT-CTATCGTT_R1<br>D1A34_12739AAD_CGTTGGTT-CTATCGTT_R2 | 121230751 | 114804516 | 105255647 | 91,68% | 26912 |
| D1AScr | Scramble antagomiR | #1 | D1AScr_11509AAD_ATCTTGGT-AACGCGGT_R1<br>D1AScr_11509AAD_ATCTTGGT-AACGCGGT_R2 | 71207839 | 62715881 | 57010006 | 90,90% | 25365 |
| D2A34 | miR-34c antagomiR | #1 | D2A34_11510AAD_AGGCCAAG-GATCTACG_R1<br>D2A34_11510AAD_AGGCCAAG-GATCTACG_R2 | 75858222 | 67866340 | 61903691 | 91,21% | 25703 |
| D2AScr | Scramble antagomiR | #2 | D2AScr_11511AAD_GCTAACGC-GCTGAACC_R1<br>D2AScr_11511AAD_GCTAACGC-GCTGAACC_R2 | 66962493 | 58611309 | 53597989 | 91,45% | 25220 |
| D3A34 | miR-34c antagomiR | #2 | D3A34_11512AAD_TCAGATAC-CGACGTTA_R1<br>D3A34_11512AAD_TCAGATAC-CGACGTTA_R2 | 65177637 | 56456916 | 51841800 | 91,83% | 25926 |
| D3AScr | Scramble antagomiR | #2 | D3AScr_11513AAD_TAACGCCA-TGGATCAA_R1<br>D3AScr_11513AAD_TAACGCCA-TGGATCAA_R2 | 78925330 | 69977368 | 64169282 | 91,70% | 26617 |
